# A novel polymerase III promoter for gene editing in the agricultural pest *Ceratitis capitata*

**DOI:** 10.64898/2026.04.21.719894

**Authors:** Amber S. K. Hall, Samuel Matthew Shackleton-Chavez, Tracey Chapman, Philip T. Leftwich

## Abstract

We report the identification and functional validation of a 7SK RNA polymerase III promoter in the Mediterranean fruit fly, *Ceratitis capitata*. CRISPR/Cas9-based genetic control strategies for this global agricultural pest, including gene drives and precision guided sterile insect approaches, require efficient guide RNA expression, yet only a single U6 Pol III promoter had previously been validated for this purpose in *C. capitata*, and no 7SK promoter had been characterised in any Tephritid species. Using comparative genomics with *Drosophila* orthologues, we identified a previously unannotated 7SK gene in the *C. capitata* genome, confirmed its transcriptional activity by RT-PCR, and demonstrated that the cloned promoter drives functional guide RNA expression in CRISPR/Cas9-mediated knockouts of the *white* gene. Comparative analysis identified putative 7SK orthologues across the Tephritid fruit flies. The availability of this additional new Pol III promoter will enable multiplexed guide RNA strategies using distinct promoters, supporting more robust genetic control designs.

**Graphical Abstract:** 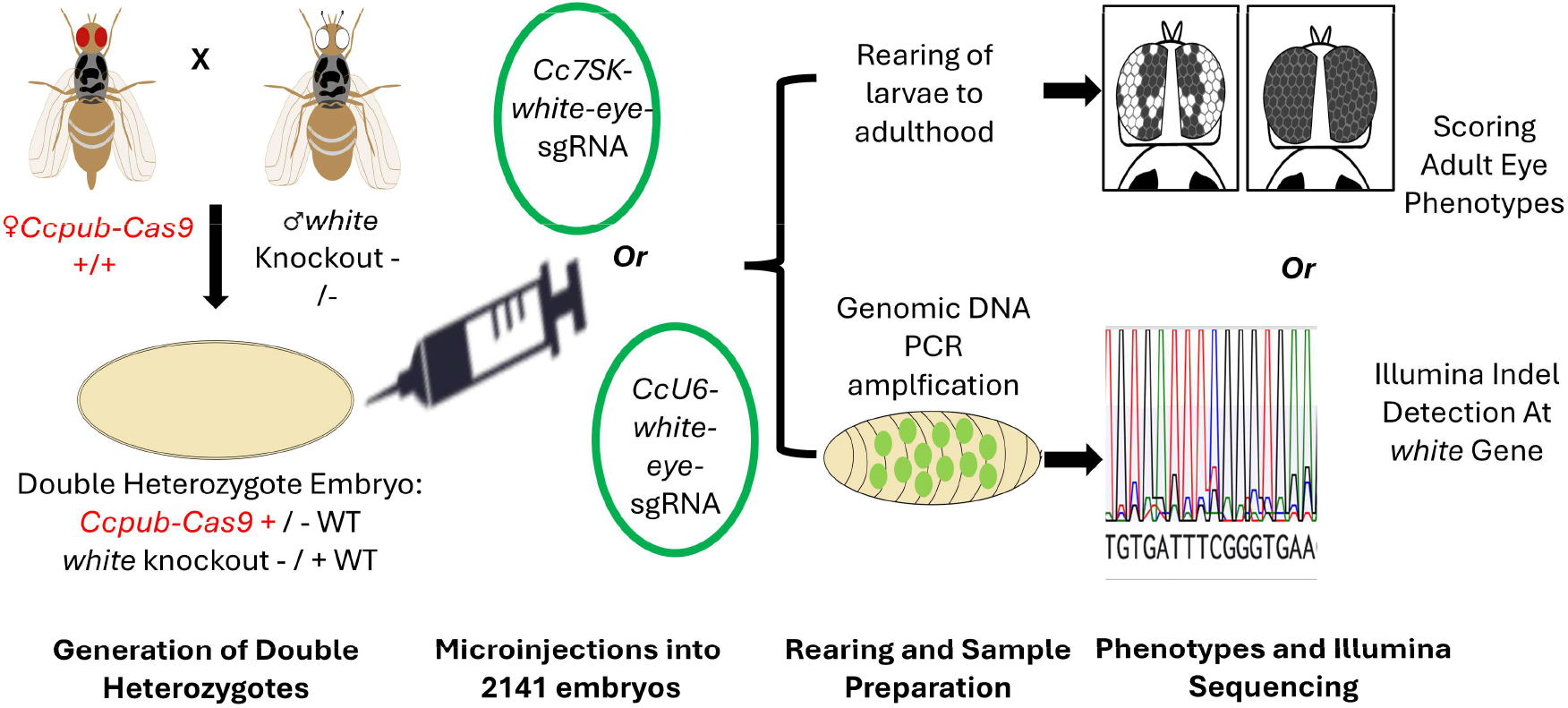

## Introduction

The Mediterranean fruit fly, *Ceratitis capitata* (medfly), is a globally significant agricultural pest within the family Tephritidae, causing damage to over 200 fruit crop species (Christenson & Foote, 1960). CRISPR/Cas9-based genetic control strategies, including homing gene drives, pgSIT, and sex conversion approaches, are under active development in this species. These strategies require the endogenous expression of two components: the Cas9 endonuclease, driven by RNA polymerase II (Pol II) promoters, and the single guide RNA (sgRNA), driven by RNA polymerase III (Pol III) promoters. Whilst the range of endonuclease promoters used in CRISPR applications is broad, gRNA promoter choice remains comparatively restricted (Verkuijl et al., 2026).

Several Pol II promoters have been evaluated for Cas9 expression in *C. capitata*, including *nanos, vasa*, and *zpg* regulatory elements. In contrast,only a single Pol III promoter, *CcU6*, has been characterised for sgRNA expression (Meccariello et al. 2021). This limitation has practical consequences. A single sgRNA target site for gene drive homing permits the accumulation of resistance alleles through non-homologous end joining repair, eventually rendering the drive inefective at the population level (Meccariello et al. 2024, Verkuijl et al. 2022). Multiplexed guide RNA designs, which target multiple sites within the same gene, represent a promising strategy to mitigate resistance (Anderson et al. 2024, Xu et al. 2025). However, expressing multiple sgRNAs from repeated copies of the same promoter increases the risk of internal recombination and cassette instability (Verkuijl et al. 2022). The use of distinct Pol III promoters for each sgRNA cassette avoids this problem and has been demonstrated successfully in mosquito gene drives (Anderson et al. 2024, Gonzalez et al. 2025).

Pol III promoters in insects are characterised by two conserved upstream elements: a proximal sequence element A (PSEA) and a TATA box (Kim et al. 2020, Hernandez et al. 2006). These motifs are conserved across Pol III promoter families, including U6 and 7SK classes, enabling identification of orthologous sequences through comparative genomics (Gruber et al. 2008).

The 7SK small nuclear RNA is a component of the 7SK ribonucleoprotein complex involved in transcriptional regulation, and its promoter has been co-opted for sgRNA expression in several insect species. In *Aedes aegypti, Anopheles stephensi*, and *Culex quinquefasciatus*, 7SK-driven sgRNA expression has proved functional for CRISPR/Cas9 editing, including in gene drive contexts (Anderson et al. 2020, Anderson et al. 2024, Gonzalez et al. 2025, Purusothaman et al. 2021).

Comparative assessments of Pol III promoters for sgRNA expression in mosquitoes support the functional utility of 7SK-class promoters. In culicine mosquito cell lines, 7SK sequences have been found to be transcriptionally active and comparable in efficiency to U6 paralogues in driving reporter expression (Anderson et al. 2020). When tested in gene drive constructs *In vivo*, 7SK promoters have yielded among the highest inheritance bias (Gonzalez et al. 2025). These findings indicate that 7SK promoters can match or exceed the activity of established U6 sequences, and that the choice of Pol III promoter can materially affect gene drive performance. No 7SK promoter has yet been identified or tested in any Tephritid species.

Here, we report the identification, validation, and functional characterisation of a 7SK Pol III promoter in *C. capitata*. We demonstrate that the *Cc7SK* promoter drives sgRNA expression sufficient to generate CRISPR/Cas9-mediated knockout of the *white* gene (chromosome 5, GeneID_101458180), establishing a second validated Pol III promoter for this species. Through comparative genomic analysis, we additionally identify putative 7SK orthologues across nine Tephritidae species, suggesting that this promoter class may be broadly available within this family of agricultural pests.

## Results & Discussion

### Identification and validation of a C. capitata 7SK RNA gene

Using the *Drosophila melanogaster* 7SK RNA sequence (FBgn0065099) as a query, we identified a putative 7SK orthologue in the *C. capitata* genome (Ccap_2.1 and EGII-3.2.1 assemblies) that had not been previously annotated. Examination of the upstream region revealed a recognisable PSEA motif and a modified TATA box bearing a cytosine substitution at the 5’ position (Figure 1). We extended this analysis across the Tephritidae, identifying a putative 7SK gene with recognisable PSEA and TATA box motifs in all nine species examined. While gene sequences were well conserved, TATA box sequences showed occasional substitutions (Figure 1, Table S1-S2)

**FIGURE 1.**
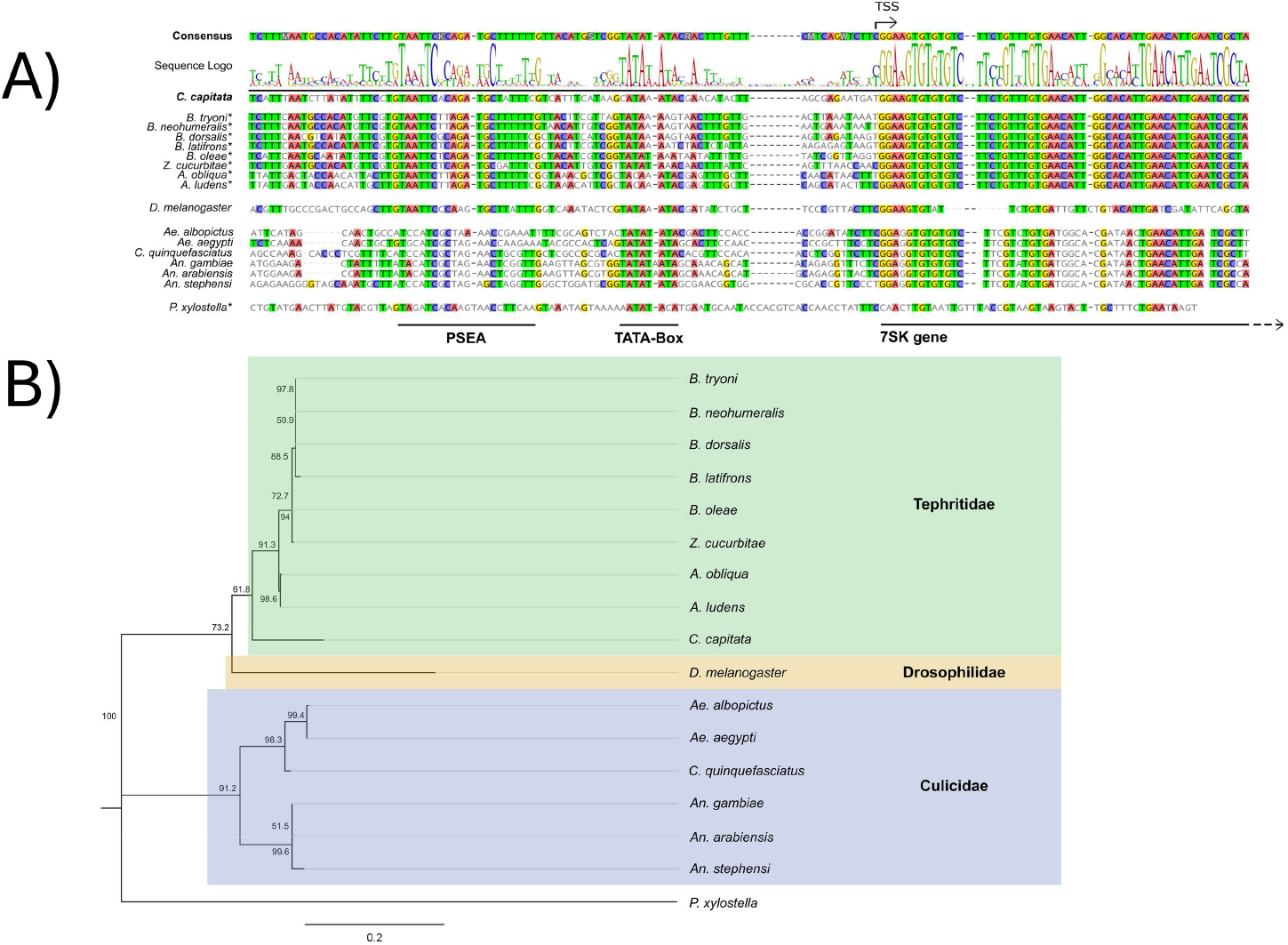
Conservation of 7SK promoter elements across dipteran species and phylogenetic relationships of 7SK genes. **(A)** Alignment of 7SK promoter sequences from *Ceratitis*, other tephritids, *Drosophila*, mosquitoes, and *Plutella*, with annotated transcription start site (TSS), proximal sequence element A (PSEA), and TATA box. **(B)** Phylogenetic relationships of 7SK genes among dipteran species, inferred by neighbour-joining under the HKY substitution model, with *Plutella xylostella* 7SK as the outgroup. Scale bar represents nucleotide substitutions per site.

To confirm that the identified locus is transcriptionally active, we performed RT-PCR on RNA extracted from adult *C. capitata* (Benakeion strain) and detected a product of the expected size (Figure S1, Table S3). This confirmed endogenous transcription from the *Cc7SK* locus.

### Generation of white knockouts using Cc7SK-driven sgRNA expression

To test whether the *Cc7SK* promoter can drive functional sgRNA expression, we cloned a 600 bp region upstream of the predicted transcription start site and used it to express a previously validated sgRNA targeting exon 3 of the *white* gene (Meccariello et al. 2017). As a positive control, we generated a parallel construct using the established CcU6 promoter with the identical sgRNA sequence. Both constructs additionally carried an AmCyan-NLS fluorescent marker to allow identification of successfully injected individuals.

Embryos for injection were generated by crossing OAMS21-Ccpub-Cas9 females (Davydova et al. 2025), which provide both integrated and maternally deposited Cas9, to homozygous white-eye mutant males. The resulting embryos were therefore heterozygous at the *white* locus, carrying one functional and one disrupted allele, such that monoallelic disruption was required for a visible phenotype change.

We injected embryos with either Cc7SK-white-eye-sgRNA (n = 584) or CcU6-white-eye-sgRNA (n = 664) plasmids. Larvae were screened at 24–72 hours post-injection for AmCyan fluorescence and separated by fluorescence score. Among adults that eclosed, we identified 9/73 (12%) mosaic white-eyed individuals from Cc7SK-injected embryos and 6 /48 (12.5%) from CcU6-injected embryos (full data in Table S4). Mosaic individuals displayed clusters of depigmented (white) ommatidia against the wild-type dark eye background, consistent with somatic disruption of *white* in a subset of cells. The recovery of mosaic phenotypes from Cc7SK-injected embryos provided direct evidence that the Cc7SK promoter drives functional sgRNA expression sufficient for Cas9-mediated cleavage in vivo (Figure 2).

**FIGURE 2.**
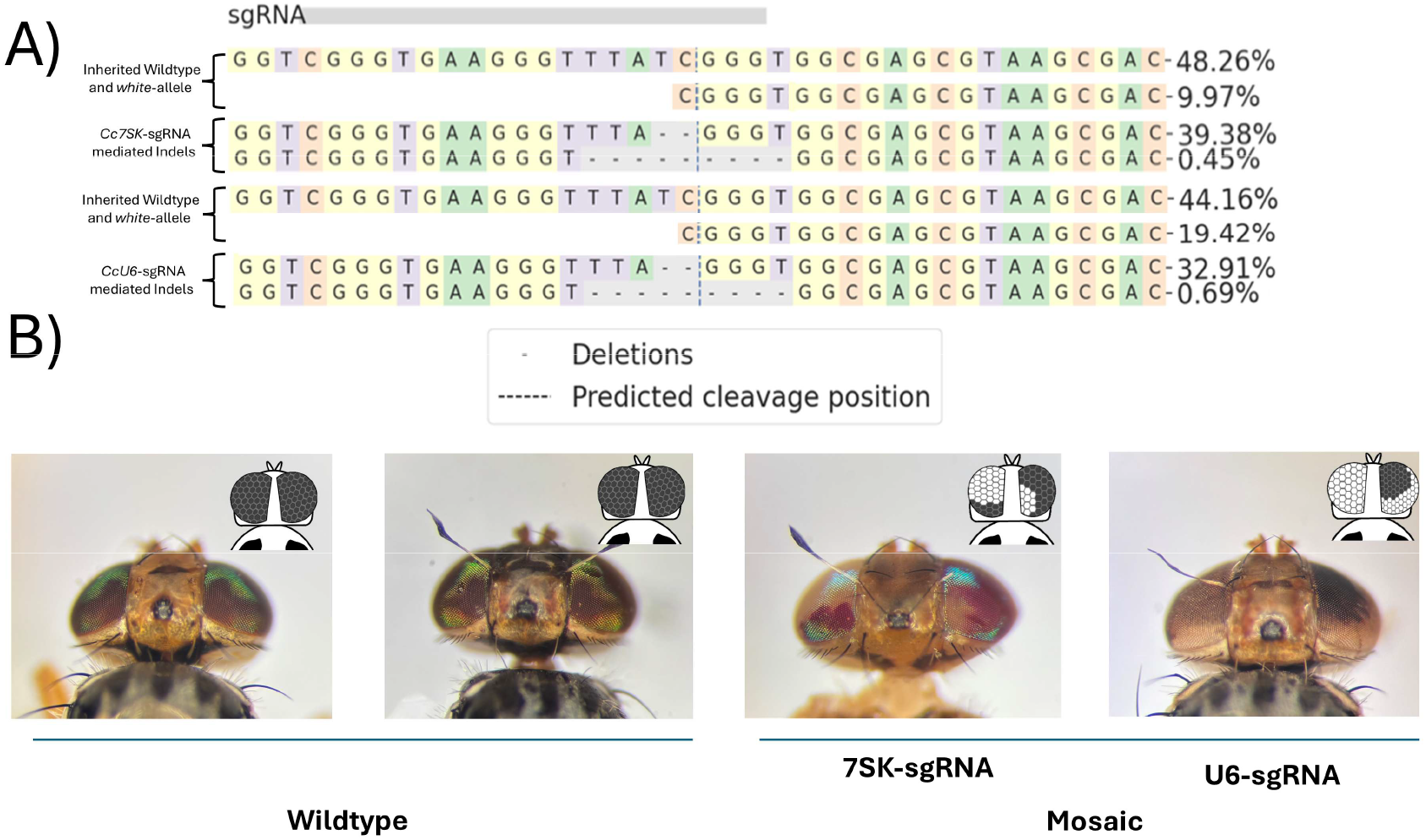
CRISPR-mediated editing of the *white* locus and resulting adult phenotypes. (**A)** Alignment of the engineered sgRNA to the *white* reference sequence (GeneID: 101458180); the grey bar indicates the sgRNA target region. Values denote the percentage of sequencing reads assigned to each sequence variant for 7SK-driven and U6-driven sgRNA constructs, respectively. The upper rows show confirmed wildtype and white mutant allele sequences; the lower rows show indel sequences identified by CRISPResso2 analysis. Source data are deposited under ENA project PRJEB111759. **(B)** Representative images of post-microinjection adult flies: wildtype (left) and mosaic mutant (right).

### Molecular confirmation of editing at the white locus

To obtain molecular evidence of target site editing, we performed a separate injection experiment using the same constructs. Embryos injected with Cc7SK-white-eye-sgRNA (n = 435) or CcU6-white-eye-sgRNA (n = 458) were screened as L1 hatchlings for AmCyan fluorescence, reared to the L2 stage, and collected for genomic DNA extraction. Fluorescence-positive larvae were pooled by construct (Cc7SK, n = 20; CcU6, n = 19) and a 258 bp amplicon spanning the sgRNA target site was sequenced using Illumina MiSeq (Amplicon-EZ, Genewiz).

Analysis with CRISPResso2 (Clement et al. 2019) revealed indels at the predicted Cas9 cleavage site in both samples (Figure 2). The predominant editing outcome under both promoters was a 2 bp deletion immediately 5’ of the PAM sequence (7SK: 39.38% and U6 32.91%). The *white* mutant allele, which carries a deletion spanning the majority of the sgRNA recognition sequence, showed minimal editing, consistent with the loss of the target site in that allele. The concordance of the dominant indel signature between both promoter constructs provides molecular confirmation that the Cc7SK promoter drives production of the same functional sgRNA as CcU6.

### Implications for genetic control construct design

These data establish Cc7SK as the second validated Pol III promoter in *C. capitata*. It is important to note that this proof-of-concept experiment, using transient plasmid-based expression, does not allow quantitative comparison of transcriptional activity between Cc7SK and CcU6. Differences in plasmid uptake during microinjection preclude meaningful assessment of relative expression levels. Previous work in *Anopheles stephensi* demonstrated that 7SK promoters can equal or exceed U6 activity for sgRNA expression when integrated and directly compared (Gonzalez et al. 2025). A comparable integrated comparison in *C. capitata*, measuring cutting or homing rates from the same genomic locus under identical Cas9 regulation, would be needed to determine the relative activity of CcU6 and Cc7SK in the germline.

The availability of a second Pol III promoter has immediate practical implications. Multiplexed gene drive designs in *Aedes aegypti* have used distinct Pol III promoters (U6 and 7SK) for each sgRNA cassette to avoid repetitive sequences that promote recombination (Anderson et al. 2024). A similar approach is now feasible in *C. capitata*, which could improve the durability of homing-based gene drives by targeting multiple sites within a gene such as transformer (Meccariello et al. 2024). Additionally, different Pol III promoters may have distinct spatiotemporal expression profiles. Having multiple promoter options therefore provides flexibility to optimise sgRNA expression timing relative to Cas9 activity in the germline.

The identification of putative 7SK orthologues in eight further Tephritidae species (Table S1) suggests that this approach may be transferable to related pest species for which genetic control tools are currently lacking.

## Methods

A putative 7SK small nuclear RNA gene was identified in the *C. capitata* genome (Ccap_2.1 and EGII-3.2.1 assemblies) by BLASTN using the *D. melanogaster* 7SK RNA sequence as a query. Upstream promoter elements (PSEA, TATA box) were identified computationally, and orthologous sequences were searched across nine Tephritidae species (Supporting Information S1-S2). Endogenous transcription was confirmed by RT-PCR on cDNA synthesised from adult Benakeion RNA (Supporting Information S3–S7).

A 600 bp region upstream of the Cc7SK transcription start site was amplified from Benakeion genomic DNA and cloned into an sgRNA expression plasmid carrying an AmCyan marker. A parallel construct using the previously characterised CcU6 promoter (Meccariello et al. 2021) with the identical white-eye sgRNA (Meccariello et al. 2017) served as a positive control. Both plasmids were verified by Nanopore sequencing (Supporting Information S8).

Embryos from crosses of OAMS21-Ccpub-Cas9 females (Davydova et al. 2025) to homozygous white-eye mutant males were injected with either construct at 500 ng/µl within 1–2 hours of oviposition. For phenotypic assessment, larvae were reared to adulthood and scored for eye colour mosaicism. For molecular confirmation, fluorescence-positive L2 larvae were pooled by construct and a 258 bp amplicon spanning the target site was sequenced on an Illumina MiSeq platform and analysed with CRISPResso2 (Clement et al. 2019). Full protocols are provided in Supporting Information S9–S16.

## Data Availability

Raw sequencing data are available through the European Nucleotide Archive under project accession PRJEB111759. Scripts and raw data can be found at (https://github.com/Philip-Leftwich/A-novel-polymerase-3-promoter-for-gene-editing-in-the-agricultural-pest-Ceratitis-capitata). Complete information on analyses can be found in Supporting Information.

## Acknowledgements

ASKH and SMSC supported by a UKRI BBSRC Norwich Research Park Biosciences Doctoral Training Partnership (Grant No. BB/T008717/1 awarded to PTL and TC). We thank Angela Meccariello and Jean-Charles de Coriolis for their advice and comments.

