## Supplemental Methods for "A novel polymerase III promoter for gene editing in the agricultural pest *Ceratitis capitata*" (from another version of this preprint: version 1, 2026-04-21)

### Supplementary Methods

#### S1. In silico identification of RNA Pol III promoters

Putative U6 and 7SK small nuclear RNA genes were identified in Tephritidae species by BLASTN (Zhang et al. 2000) searches of publicly available genome assemblies retrieved from NCBI. The *Drosophila melanogaster* U6:Aa snRNA gene sequence and the *D. melanogaster* 7SK1 snRNA gene sequence were used as queries against *C. capitata* (Ccap_2.1, GCA_000347755.4; EGII-3.2.1, GCA_905071925.1) and eight additional Tephritidae genome assemblies (Table S1). Each putative snRNA sequence was screened for the presence of an RNA polymerase III terminator sequence (TTTT motif). Sequences lacking this motif were excluded from further analysis.

Upstream promoter regions (~400 bp 5' of each snRNA gene) were retrieved and screened for conserved proximal sequence element A (PSEA) and TATA box motifs. An R script (R version 4.3) utilising the stringr (Wickham 2023) package was used to identify motifs and identify PSEA and TATA-box motifs within the promoter region sequence upstream of the putative 7SK genes. Table S2 depicts the sequence patterns provided for PSEAs and TATA-boxes.

#### S2. Multiple Sequence Alignment and Phylogeny

Putative gene and promoter sequences identified using the method described above, containing a PSEA and TATA-like motif were taken forward for alignment. Annotated 7SK genes and promoter sequences from species of mosquito (Culicidae) were also included for reference in alignments. Multiple sequence alignments were performed using the EBI browser version of MAFFT (Madeira, et al., 2024) with no alterations to standard parameters. Further analysis and visualisation were completed with Geneious Prime (v2026.0) to generate consensus sequence and sequence logo.

Consensus phylogenetic tree was constructed and visualised with Geneious Prime (v2026.0). 7SK gene sequences were aligned using MAFFT, excluding any regulatory regions 3’ and 5’ of the gene sequences. Relationships were inferred using neighbour-joining with genetic distances calculated under HKY substitution model. Branch support assessed using bootstrap of 1000 replicates. *P. xylostella* was included as an outgroup.

#### S3. Genomic DNA extraction

Genomic DNA was extracted from *C. capitata* (Benakeion strain) using the DNeasy Blood and Tissue Kit (Qiagen, Cat. No. 69504) with the following modifications. Flies were stored at −20 °C for at least 24 hours prior to extraction. Samples of 1–5 flies were ground with a pestle in a 1.5 ml microcentrifuge tube before addition of 180 µl Buffer ATL and 20 µl Proteinase K. Samples were vortexed briefly and incubated at 56 °C, with vortexing every 5 minutes. For standard extractions (promoter cloning, reference sequence generation), incubation was 16 minutes. For L2 larval gDNA extraction used in amplicon sequencing, larvae were pooled in batches of 5 (four batches per construct; one CcU6 batch contained 4 larvae) and incubated at 56 °C after addition of Buffer AL. Following incubation, 200 µl Buffer AL was added and samples were incubated for a further 10 minutes. 200 µl of 100% ethanol was added and mixed thoroughly by vortex. Samples were centrifuged briefly and the supernatant was transferred to spin columns, avoiding transfer of incompletely digested tissue. Wash steps with Buffers AW1 and AW2 were carried out according to the manufacturer's protocol. DNA was eluted with 200 µl Buffer AE, incubated for 1 minute at room temperature, and centrifuged for 1 minute at maximum speed. DNA concentration and purity were assessed by spectrophotometric analysis (NanoDrop 2000, Thermo Scientific).

#### S4. RNA extraction, DNase treatment, and cDNA synthesis

Total RNA was extracted from adult *C. capitata* tissues. Where residual genomic DNA contamination was detected, RNA samples were treated with TURBO DNase (TURBO DNA-free Kit, Invitrogen) following the routine DNase treatment protocol. RNA was diluted to approximately 100 µg/ml and a 25 µl reaction was prepared with 2.5 µl 10× TURBO DNase Buffer and 1 µl TURBO DNase enzyme, then incubated at 37 °C for 20 minutes. DNase Inactivation Reagent (2.5 µl) was vortexed to resuspend, added to each sample, and incubated at room temperature for 5 minutes with periodic mixing. Samples were centrifuged at 10,000 × g for 1.5 minutes and the supernatant was transferred to a clean tube.

cDNA was synthesised using the QuantiTect Reverse Transcription Kit (Qiagen) following the quick-start protocol. RNA samples were first treated with gDNA Wipeout: 11 µl RNase-free water, 2 µl gDNA Wipeout Buffer, and 1 µl template RNA were combined, incubated at 42 °C for 2 minutes, and placed immediately on ice. The reverse transcription reaction was prepared on ice: 1 µl Quantiscript Reverse Transcriptase, 4 µl Quantiscript RT Buffer (5×), 1 µl RT primer mix, and 14 µl of the gDNA elimination reaction. The reaction was incubated at 42 °C for 15 minutes, followed by 95 °C for 3 minutes, then held at 4 °C. cDNA concentration and purity were measured using a NanoDrop 2000.

#### S5. RT-PCR validation of endogenous 7SK transcription

RT-PCR was performed on cDNA (S3) to confirm transcriptional activity at the putative Cc7SK locus. Primers were designed against the identified 7SK RNA gene sequence (Table S3). PCR was performed using DreamTaq PCR Master Mix (2×) with the following conditions: initial denaturation at 95 °C for 2 minutes; 35 cycles of 95 °C for 30 seconds, 51–53 °C for 30 seconds, and 72 °C for 1 minute; final extension at 72 °C for 15 minutes; hold at 18 °C. gDNA and no-template controls were included. Products were analysed by agarose gel electrophoresis (1% agarose, 1× TBE buffer). RT-PCR gel images are presented in Figure S1.

#### S6. PCR protocols and primer design

Primers were designed manually and using the Primer 3 (Untergasser 2012) primer wizard against identified U6 and 7SK gene and promoter sequences (Table S3). Annealing temperatures were calculated using the NEB Tm Calculator (https://tmcalculator.neb.com). Primers were synthesised by Merck and diluted to a working concentration of 100 µM.

Standard PCR reactions contained: 12.5 µl DreamTaq PCR Master Mix (2×), 1.25 µl forward primer (10 µM), 1.25 µl reverse primer (10 µM), and approximately 1 µg template DNA, made up to 25 µl with nuclease-free water. Cycling conditions were: 95 °C for 2 minutes; 35 cycles of 95 °C for 30 seconds, 51–53 °C for 30 seconds (adjusted per primer pair), 72 °C for 1 minute; 72 °C for 15 minutes; hold at 18 °C.

For high-fidelity applications (promoter amplification, amplicon generation for sequencing), Q5 High-Fidelity 2× Master Mix (NEB, Cat. No. M0492S) was used according to the manufacturer's protocol, with annealing temperatures adjusted per primer pair (63 °C for amplicon sequencing primers).

#### S7. Agarose gel electrophoresis

PCR products were analysed by gel electrophoresis on 1% agarose gels in 1× TBE buffer unless otherwise stated.

#### S8. Cloning of sgRNA expression plasmids

Two sgRNA expression plasmids were constructed: CcU6-white-eye-sgRNA and Cc7SK-white-eye-sgRNA. Both expressed the same white-eye sgRNA sequence previously described in [REF: Meccariello et al. (2017)] and carried an AmCyan fluorescent marker. The plasmids were identical except for the Pol III promoter sequence.

The Cc7SK promoter sequence (600 bp upstream of the predicted transcription start site) was amplified from Benakeion strain gDNA using Q5 High-Fidelity 2× Master Mix. The Cc7SK-white-eye-sgRNA construct was assembled by Gibson assembly: the linearised vector backbone was incubated with the amplified Cc7SK promoter insert, followed by heat-shock transformation into chemically competent cells. Cells were supplemented with SOC medium and incubated at 37 °C for 60 minutes in a shaking incubator before plating onto selective plates and overnight incubation at 37 °C. Colonies were picked, inoculated into LB medium with the appropriate antibiotic, grown overnight, and plasmid DNA was recovered by miniprep. The CcU6-white-eye-sgRNA construct was assembled by Gibson and restriction-ligation assemblies.

Both plasmids were purified by midiprep (NucleoBond Xtra Midi EF, Macherey-Nagel, Cat. No. 740420.10) and further purified using Ultrafree centrifugal filters (Millipore, Cat. No. UFC30GV0S). Plasmid sequences were verified by Nanopore sequencing (Plasmidsaurus). Complete plasmid sequences are available from the European Nucleotide Archive under project accession PRJEB111759

#### S9. Medfly rearing and crosses

*C. capitata* were maintained on a 12:12 hour light:dark cycle at 25.2 °C (±1.5 °C) and 65% (±5%) relative humidity. Adults were provided with a 3:1 sucrose:hydrolysed yeast mixture and water ad libitum from moistened cotton.

Embryos for injection were generated by crossing homozygous OAMS21-Ccpub-Cas9 females (Davydova et al. 2025) to homozygous *white* mutant males (carrying a deletion: GCGTGC-[21 bp deletion]-CGGGTG, generated by sgRNA CGGGTGAAGGGTTTATCGGGTGG). Crosses were established as 30 virgin females to 10 males in 10 × 10 × 10 cm cages with fine fabric mesh floors suspended above distilled water for egg collection. Adults were mated from day 0 post-eclosion; eggs were collected for injection from days 4–10 post-eclosion. Transgenic and mutant lines were kindly provided by Angela Meccariello.

#### S10. Embryo microinjection

Embryos were collected 1–2 hours post-oviposition and dechorionated manually using a hypodermic needle. Embryos were aligned anterior-end towards the injection needle on glass coverslips (22 × 22 mm, No. 2) coated with a thin layer of n-heptane (Fluka, Cat. No. 34495) mixed with double-sided adhesive tape. Embryos were desiccated for 6 minutes in sealed Petri dishes with granular calcium chloride (Fluka, Cat. No. C1016), then coated with Halocarbon Oil 700 (Sigma, Cat. No. H8898).

Embryos were collected in batches of 3–6 slides (mean 22.4 embryos per slide for CcU6, 21.2 for Cc7SK). Injections were performed immediately using hand-pulled quartz needles on an Eppendorf FemtoJet 4i microinjector (Cat. No. E5252000021). CcU6-white-eye-sgRNA and Cc7SK-white-eye-sgRNA plasmids were diluted to 500 ng/µl in molecular-grade water and kept at 4 °C. For each injection, 2–3 µl was loaded into the needle. A continuous compensation pressure of 32–237 hPa was applied, with the needle inserted 10–20% of the embryo body length into the anterior pole and held for 1–2 seconds until a visible bolus was observed. Injected embryos were returned to standard rearing conditions.

#### S11. Fluorescence screening and larval sorting

Larvae were screened for AmCyan fluorescence at 24-, 48-, and 72-hours post-injection using a fluorescence microscope (Leica, Type 11307072057) fitted with a GFP Plus filter and an HBO 103 W/2 mercury lamp (Osram).

Larvae were scored and separated into two groups. Fluorescence-positive larvae were defined as alive with green fluorescence, including a minority with visible nuclear localisation signal. Fluorescence-negative larvae were defined as those showing no fluorescence, red/orange bleed-through from the parental OAMS21-Ccpub-Cas9 transgene, autofluorescence only, or dead.

#### S12. Larval rearing

*Rearing to adulthood:* Fluorescence-scored L1 larvae were transferred by hypodermic needle onto carrot-based media in Petri dishes (S13). On day 5 post-injection, dishes were placed uncovered into sealed pupation boxes containing a layer of sand with ventilation holes. Pupae were collected on day 14 and allowed to eclose in sealed Petri dishes, typically between days 19 and 22 post-injection.

*Rearing to L2 stage for sequencing:* L1 larvae were transferred individually into wells of a 48-well plate pre-loaded with approximately 2–3 g of carrot-based media. Larvae were reared for a further 48–72 hours (up to day 6 post-injection), re-scored for fluorescence, then transferred to 1.5 ml microcentrifuge tubes and stored at −80 °C.

#### S13. Larval media preparation

Carrot-based larval media was prepared as follows: 900 g fresh carrots, 11.4 g sodium benzoate (Thermo, Cat. No. A15946.30), 16.4 ml hydrochloric acid (32%, Sigma-Aldrich, Cat. No. 1.00313.2500), 2.2 litres reverse osmosis water, 120 g yeast (MP Biomedicals, Cat. No. 903312), and 357 g carrot powder (BuyWholeFoodsOnline, Cat. No. CONFSKU946988). Protocol kindly provided by Angela Meccariello.

#### S14. Adult eye phenotype imaging

Injection survivors that eclosed (days 19–22 post-injection) were scored for eye phenotype at 4 days post-eclosion using a stereo microscope. Eye phenotypes were imaged post-mortem using a Leica MZ75 stereo microscope with a Leica CLS 150X light source.

#### S15. Generation of reference *white* sequences

Wildtype and mutant-allele reference sequences for the *white* locus were generated for comparison with amplicon sequencing data.

For the Benakeion wildtype sequence, gDNA was extracted from individual flies (S3 and amplified using primers white_2.3kb_F and white_2.3kb_R (Table S1) with Q5 High-Fidelity 2× Master Mix. The amplicon was sequenced by linear/PCR amplicon sequencing (Plasmidsaurus).

For the *white* mutant-allele sequence, gDNA was amplified using primers F_indel_W and R_indel_W (Table S3), derived from Meccariello et al. (2024). Sanger sequencing was performed by Eurofins Genomics (TubeSeq Supreme) using the forward primer F_indel_W.

#### S16. Amplicon sequencing and CRISPResso2 analysis

L2 larvae collected at −80 °C (S12) were pooled by construct for gDNA extraction. PCR targeting the *white* editing locus was performed using primers F_indel_Wi and R_indel_Wi (Table S3), which are modified versions of F_indel_W and R_indel_W with Illumina partial adapter overhangs. Reactions used Q5 High-Fidelity 2× Master Mix with an annealing temperature of 63 °C. Amplicon size was verified by agarose gel electrophoresis (expected: 323 bp, comprising 258 bp target amplicon plus 65 bp adapter sequence). Amplicons were purified using the Monarch PCR and DNA Cleanup Kit (NEB, Cat. No. T1130S).

Purified amplicons were pooled by construct (four batches each), normalised by molarity, and submitted for paired-end sequencing (Amplicon-EZ, Genewiz) on an Illumina MiSeq platform. FASTQ files were analysed using CRISPResso2 (Clement et al. 2019). Analysis scripts are available at <https://github.com/Philip-Leftwich/A-novel-polymerase-3-promoter-for-gene-editing-in-the-agricultural-pest-Ceratitis-capitata>.

### Supplementary Figures

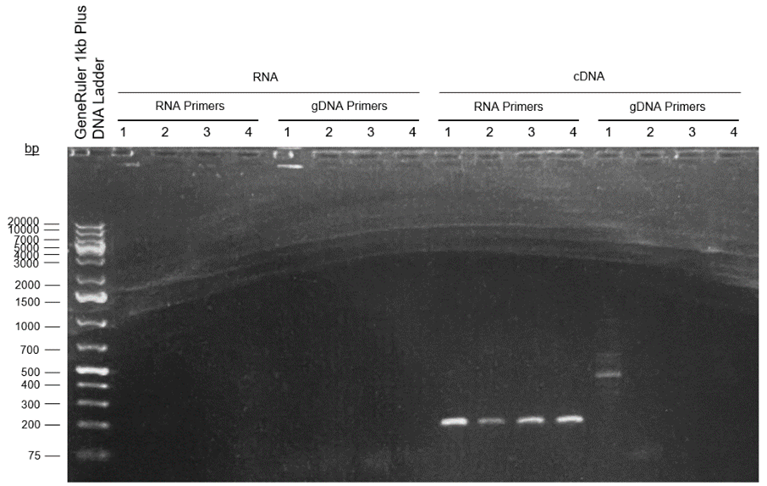

**Figure S1.** Amplification of cDNA with gDNA primers and RNA primers. PCRs were run on RNA post gDNA wipeout step of Reverse Transcriptase (RT) cDNA synthesis and on newly synthesised cDNA. Lanes refer to RNA sample: 1 - female ovaries; 2 - female soma; 3 - male testes; 4 - male abdomen. Successful amplification of the 7SK RNA region fragments from cDNA using RNA primers yielded bands of ~211bp. Partial gDNA contamination present in cDNA sample 1-5.

### Supplementary Tables

**Table S1*.*** Accession codes and genome assemblies for 7SK genes and promoters for mosquito species and putative Tephritidae and *P. xylostella* 7SK sequences retrieved from BLASTN analysis with reference to genome assembly and sequence retrieved from NCBI.

| **Species** | **Database** | **Genome Assembly** | **Sequence Accession** | **Previously Annotated** |
| --- | --- | --- | --- | --- |
| Ceratitis capitata | NCBI | EGII-3.2.1 | CAJHJT010000012.1 | No |
| Anastrepha Ludens | NCBI | idAnaLude1.1 | NC_071501.1 | No |
| Anastrepha obliqua | NCBI | idAnaObli1_1.0 | NC_072896.1 | No |
| Bactrocera dorsalis | NCBI | ASM2337382v1 | NC_064303.1 | No |
| Bactrocera neohumeralis | NCBI | APGP_CSIRO_Bneo_wtdbg2-racon-allhic-juicebox | NC_065920.1 | No |
| Bactrocera tryoni | NCBI | CSIRO_BtryS06_freeze2 | NC_052500.1 | No |
| Bactrocera latifrons | NCBI | ASM185335v1 | NW_017536630.1 | No |
| Bacterocera oleae | NCBI | idBacOlea1 | NC_091537.1 | No |
| Zeugodacus cucurbitae | NCBI | idZeuCucr1.2 | NC_071668.1 | No |
| Drosophila melanogaster | FlyBase | Release 6 plus ISO1 MT (GCF_000001215.4) | FBgn0065099 | Yes |
| Aedes aegypti | VectorBase | AaegL5_3 | AAEL018514 | Yes |
| Aedes albopictus | VectorBase | JXUM01S075709 | AALF029648 | Yes |
| Culex quinquefasciatus | VectorBase | DS232527 | CPIJ039933 | Yes |
| Anopheles arabiensis | VectorBase | KB704651 | AARA015292 | Yes |
| Anopheles gambiae | VectorBase | AgamP4_3L | AGAP028235 | Yes |
| Anopheles stephensi | VectorBase | KE388938 | ASTEI12173 | Yes |
| Plutella xylostella | NCBI | ilPluXylo3.1 | NC_063983.1 | No |

**Table S2** – 7SK promoter sequence patterns for TATA-like and PSEA elements were devised from *Drosophila* 7SK motifs, previously described by (Gruber et al., 2008). Position 5’ of gene refers to number of base pairs upstream of TSS. Sequences would be returned if either or both a TATA-like and PSEA motif were identified within the given positions alike in the promoter region sequence

| **Promoter Element Motifs** | **Position 5’ of TSS** | **Sequence Pattern** |
| --- | --- | --- |
| TATA-like | Between 22bp and 31bp | (A\|C\|T)(A\|T)TA(A\|T)A |
| PSEA | Between 40bp and 50bp | TAATTCCCAAGTGCTTATTTG |

**Table S3*.*** Primers used for amplicon generating, sequencing and reference sequence generation.

| **Primer Name** | **Sequence (5’ to 3’)** |
| --- | --- |
| RNA-Fwd (Fwd7SK1-3) | GACGGTCACACTCTCCAGTCAA |
| RNA-Rev (Rev7SK1-3) | GAAATGACGTCGCAGAGGAACA |
| gDNA-Fwd (Fwd7SK1-2) | ACCTGCTTATTGCTGCTTTGCA |
| gDNA-Rev (Rev7SK1-2) | ACCCACAAACAACACCACCAGCA |
| F_indel_Wi | ACACTCTTTCCCTACACGACGCTCTTCCGATCTACGGCATATGACACAAAAACAG |
| R_indel_Wi | GACTGGAGTTCAGACGTGTGCTCTTCCGATCTTTCTGCGATAGCTTTTTCAACA |
| F_indel_W | ACGGCATATGACACAAAAACAG |
| R_indel_W | TTCTGCGATAGCTTTTTCAACA |
| white_2.3kb_F | CAGTTGCTCTTTCTACTCCGAAATTG |
| white_2.3kb_R | GAAACATTGGCTACTAAAGTGACA |

**Table S4*.*** Total Microinjection numbers, hatching rates and fluorescence numbers, adult eclosions and mosaicisms

| **Plasmid** | **Injections** | **Hatchlings Total (Fluorescent/Non-Fluorescent)** | **Adults**  **(Wildtype/Mosaic)** |
| --- | --- | --- | --- |
| cc-U6-gRNA-white-eye | 664 | 109  (78/31) | 48  (42/6) |
|  | 458 | 73  (43/30) | NA |
| cc-7SK-gRNA-white-eye | 584 | 131  (99/32) | 73  (64/9) |
|  | 435 | 59  (39/20) | NA |
